# Recurrent co-option and recombination of cytokine and three finger proteins in multiple reproductive tissues throughout salamander evolution

**DOI:** 10.1101/2022.01.04.475003

**Authors:** Damien B. Wilburn, Christy L. Kunkel, Richard C. Feldhoff, Pamela W. Feldhoff, Brian C. Searle

## Abstract

The proteomic composition of amphibian gametes is largely a molecular mystery, particularly for Urodeles (salamanders and newts) which have few genomic-scale resources. Lungless salamanders (family Plethodontidae) include approximately two thirds of all extant salamander species and are classic models of vertebrate mating behavior. As part of an extended, multi-stage courtship ritual, male plethodontid salamanders deliver rapidly evolving protein pheromones that modify female behavior and improve male reproductive success. Despite great interest in this set of pre-mating reproductive barriers, limited characterization of plethodontid gametes has prohibited investigation of post-mating pre-zygotic barriers such as sperm-egg recognition. In this study, we performed transcriptomic analyses of testis and ovary using long-read PacBio sequencing and proteomic analyses of sperm using mass spectrometry for two evolutionary divergent plethodontid species, *Plethodon shermani* and *Desmognathus ocoee*. In both species, many of the most abundant sperm proteins were paralogs of the courtship pheromones Plethodontid Receptivity Factor (PRF), Plethodontid Modulating Factor (PMF), and Sodefrin Precursor-like Factor (SPF). Sperm-specific paralogs of PMF and SPF are likely the most abundant secreted proteins in *P. shermani* and *D. ocoee*, respectively. In contrast, sperm PRF lacks a signal peptide and may be expressed in cytoplasm. PRF pheromone genes evolved independently multiple times through repeated gene duplication of sperm PRF genes and signal peptides recovered by recombination with PMF genes. Phylogenetic analysis of courtship pheromones and their sperm paralogs support that each protein family evolved for these two reproductive contexts at distinct evolutionary time points between 17 and 360 million years ago. As the first molecular characterization of salamander gametes, this study expands our knowledge of amphibian fertilization beyond frogs and provides novel insight into the evolutionary processes by which new, rapidly evolving reproductive proteins may evolve.

## Introduction

Sexual reproduction is a hallmark of most animal life cycles and is the process by which progeny are generated from the recombination of two genomes, usually from individuals of the same species. Numerous types of reproductive barriers exist to restrict such recombination – including physical boundaries, niche utilization, asynchronous mating cycles, and genetic incompatibilities – and the formation of such barriers is what ultimately transforms interbreeding populations into discrete species (Rundle and Nosil, 2005). While ecological and genetic forces have been well investigated as drivers of speciation, the role of molecular prezygotic barriers such as gamete recognition on speciation has been a topic of increasing relevance in diverse eukaryotic taxa, including fungi (Jones and Bennett, 2011), plants (Pease et al., 2016), insects (Swanson et al., 2001;Noh and Marshall, 2016), mollusks (Wilburn et al., 2018), fish (Herberg et al., 2018), and mammals (Monne et al., 2008;Avella et al., 2014). In nearly all eukaryotic species, the genes associated with reproduction are among the fastest evolving in their respective genomes, rivaling or exceeding the rate of immune gene evolution (Swanson and Vacquier, 2002;Wilburn and Swanson, 2016). In contrast to genes associated with somatic phenotypes, reproduction-related genes can evolve through a runaway sexual selection process that can yield unparalleled rates of molecular evolution and are hypothesized to contribute to speciation (Arnold and Houck, 2016;Wilburn and Swanson, 2016;Wilburn et al., 2017). However, explicitly testing such hypotheses has been hampered by our limited understanding of the biochemical mechanisms underlying sexual reproduction, including sperm-egg interactions in animals (Wilburn and Swanson, 2016). Recently published structural and biochemical studies have begun to provide insight into the molecular basis of egg-sperm recognition in marine invertebrates, fishes, birds, and mammals (Ichikawa et al., 2016;Raj et al., 2017;Herberg et al., 2018;Wilburn et al., 2018), but among vertebrates substantially less attention has been given to reptiles and amphibians. This is especially true for salamanders, where to date there have been no studies characterizing the molecular composition of salamander gametes. This is largely consistent with a general lack of molecular resources for salamanders, whose genomes are ~5-30X larger than humans (depending on the species) and have been immensely difficult to assemble using short-read sequencing technologies (Nowoshilow et al., 2018;Sessions and Wake, 2021). In this study, we present the first molecular characterization of salamander gametes for two species that are established models of reproduction and prezygotic mating barriers.

Lungless salamanders (family Plethodontidae) are the most speciose family of salamanders (496 out of 766 species) with expansive radiations throughout the North American coasts and central America that are classic models of mating behavior and pheromone communication (Shen et al., 2015;AmphibiaWeb, 2021). Male and female salamanders engage in a highly orchestrated courtship ritual whose precise details vary by species, but a common feature to all plethodontid salamanders is the inclusion of a “tail straddling walk.” During this period, the female salamander will straddle the tail of the male while the pair walk in unison for up to an hour, and this prepares the female to receive a spermatophore that the male will deposit on the substrate (Houck and Arnold, 2003). In most plethodontid species, male salamanders deliver non-volatile protein pheromones to females during the tail straddling walk via a chin gland (referred to as the mental gland) that can alter her mating behavior, including reducing the length of the walk (Wilburn et al., 2017). The males of most species transdermally deliver pheromones by abrading the female dorsum with skin protrusions called “premaxillary teeth” (Houck and Arnold, 2003). However, in a single clade of *Plethodon* species restricted to eastern North America, males possess a large pad-like mental gland that is “slapped” against the female snout, delivering pheromones to the olfactory system, activating neurons that project to the brain, and directly alter female neurophysiology and mating behavior (Rollmann et al., 1999;Wirsig-Wiechmann et al., 2002;Laberge et al., 2008;Wilburn and Feldhoff, 2019). The slapping and scratching pheromone delivery systems have been most thoroughly examined in the red-legged salamander (*Plethodon shermani*) and the Ocoee salamander (*Desmognathus ocoee*), respectively. From these two species, three major pheromone families have been chemically purified and experimentally demonstrated to alter female mating behavior: Sodefrin Precursor-like Factor (SPF), Plethodontid Modulating Factor (PMF), and Plethodontid Receptivity Factor (PRF) (Rollmann et al., 1999;Houck et al., 2007;Wilburn et al., 2015).

All three major pheromone families (SPF, PMF, and PRF) are rapidly evolving within Plethodontidae and have been co-opted for pheromone function at different evolutionary times (Watts et al., 2004;Palmer et al., 2007;Palmer et al., 2010). SPF is the oldest known pheromone family in vertebrates, with homologs shared between salamanders and anurans (frogs and toads) that have been separated by ~360 million years (Bossuyt et al., 2019). PMF is also an ancient pheromone likely found in the last common ancestor of all plethodontid salamanders (~66 million years ago) and is homologous to the three-fingered protein (TFP) superfamily (Wilburn et al., 2012;Shen et al., 2015). Despite this homology, PMF is not a classical TFP as changes in the disulfide bonding pattern have resulted in a unique topology that increases structural flexibility in rapidly evolving regions of the molecule (Wilburn et al., 2014a). SPF is also related to the TFP superfamily, but with two TFP-like domains that have even more distinct disulfide bonding patterns (Doty et al., 2016). The structural consequences of this altered disulfide pattern are unknown as a structure of SPF has not yet been solved. In contrast, PRF is a relatively new pheromone that is only found in *Plethodon* species of eastern North America (~17 million years old) and is related to IL-6 cytokines (Rollmann et al., 1999;Shen et al., 2015). In the models for slapping (*P. shermani*) and scratching (*D. ocoee*) delivery, PRF and PMF are the major pheromones in *P. shermani* versus SPF being dominant in *D. ocoee*. All three families are highly paralogous and each species usually has multiple sequence isoforms of each pheromone (Wilburn et al., 2012;Chouinard et al., 2013;Doty et al., 2016;Wilburn et al., 2017). In addition to isoform variation within each species, all three pheromone families have experienced rapid evolutionary change via positive selection. This accelerated evolution is hypothesized to be a product of coevolution with female receptors and may function to restrict mating between closely related species that share overlapping habitat ranges (Watts et al., 2004;Palmer et al., 2005;Palmer et al., 2007;Palmer et al., 2010;Wilburn et al., 2012).

While pre-mating reproductive barriers of plethodontid salamanders have been well studied, post-mating pre-zygotic barriers such as gametic recognition between egg and sperm have not yet been investigated. Little is known about the molecular composition of amphibian gametes beyond studies of a single frog genus (*Xenopus spp*.), and it is unknown if salamanders possess homologs of many common gamete-associated genes. Salamander genomes are extraordinarily large (~5-30X the human genome) which has challenged their sequencing and assembly (Nowoshilow et al., 2018). Consequently, salamanders have historically been excluded from comparative genomic analyses that would identify such genes. In this study, we provide the first detailed molecular description of salamander gametes based on long-read transcriptomic analyses of plethodontid ovary and testis, paired with proteomic analysis of plethodontid sperm from the model species *P. shermani* and *D. ocoee*. A surprising result was the discovery that sperm express high levels of paralogs to all three pheromone families. Through molecular phylogenetic analysis, we find that each family was independently co-opted at different times throughout plethodontid and amphibian evolution. Careful examination of the pheromone families, their sperm paralogs, and homologous vertebrate proteins revealed a complex evolutionary pattern where two protein families have been repeatedly co-opted for reproductive functions through recurrent gene duplications, rapid evolution, and recombination.

## Materials and Methods

### Animal care and biological sample collection

Plethodontid salamanders were collected during their breeding season in early August, with *P. shermani* collected at Wayah Bald (Macon County, North Carolina; 35°10’48” N, 83°33’38” W) and *D. ocoee* collected at Deep Gap (Clay County, North Carolina; 35°02’20” N, 83°33’08” W). Salamanders were sexed based on large ova visible through the ventral body wall in females, the presence of a large mental gland in male *P. shermani*, and premaxillary teeth in male *D. ocoee*. Animals were temporarily maintained at Highlands Biological Station where they were individually housed in clean plastic boxes (17 x 9 x 13 cm) lined with a damp paper towel and a second damp crumpled paper towel as a refuge, with temperature and humidity maintained at 15-18°C and ~70% humidity, respectively. Animals were transferred weekly to clean boxes with fresh substrate and fed two waxworms (*Galleria mellonella*). Testis and ovary samples were collected from two male and female specimens of each species by briefly anesthetizing animals in 7% diethyl ether in water, euthanizing them by rapid decapitation, quickly dissecting reproductive tissue from the body cavity, and preserving tissue samples in RNAlater (Ambion). For sperm sample collection, staged mating trials were performed as adapted from Wilburn et al. (2015). Briefly, a single male and female salamander from the same species were paired in a clean plastic box lined with a damp paper towel under limited red light. Pairs of salamanders were visually monitored for courtship behavior, and if a pair successfully performed a tail straddling walk and a spermatophore was deposited on the substrate, the courtship was disrupted to prevent insemination, and the spermatophore collected using sterile forceps. Spermatophores were gently triturated in 100 μL amphibian Ringer’s solution for 15 minutes to release sperm and soluble components from the spermatophore casing, insoluble components (including sperm cells) collected by centrifugation at 10,000 x *g* for 10 minutes, and then stored in RNAlater. All animals were collected with appropriate permits from the North Carolina Wildlife Resources Commission and animal care protocols were reviewed and approved under the Highlands Biological Station Institutional Animal Care and Use Committee (Protocol #15-07).

### RNA isolation and gonad transcriptome sequencing

RNA was isolated from individual testis and ovary samples of both *P. shermani* and *D. ocoee* using a combination of Trizol extraction and silica column purification. First, tissue samples were homogenized in 1 mL Trizol (Thermo Fisher), insoluble material removed by brief centrifugation, the sample incubated for 3 min at room temperature following addition of 0.2 mL chloroform and vortexing, centrifuged at 14,000 *x g* for 15 minutes at 4°C, and the supernatant was collected. Second, the supernatant was mixed with 0.35 mL ethanol, applied to a RNeasy column (Qiagen) with centrifugation 10,000 *x g* for 20 sec, and serially washed with 0.2 mL Buffer RW1 and twice with 0.5 mL Buffer RPE with matching centrifugation steps of 10,000 *x g* for 20 sec. The collection tube was replaced after the Buffer RW1 and following the final RPE wash, residual ethanol was removed by a final centrifugation step at 10,000 *x g* for 3 min. RNA was eluted into 50 μL RNase-free water and concentration estimated based on 260 nm absorbance. To prepare RNA for PacBio Iso-Seq analysis, single-stranded cDNA was generated from RNA using the SMARTer cDNA synthesis kit (Clontech, Palo Alto, CA) with tagged oligo-dT primers that include one of eight 15-bp barcodes to distinguish individual samples (2 species x 2 tissues x 2 biological replicates). Double stranded cDNA was prepared using Accuprime High-Fidelity *Taq* polymerase (Thermo Fisher) with PCR cycles of 95°C for 15 sec, 65°C for 30 sec, and 68°C for 6 min, preceded by a single initial melting phase of 95°C for 2 min. Cycle number was optimized for each sample to avoid overamplification. The cDNA was purified using AmpureXP beads (Pacific Biosciences) at final concentrations of both 1.0X beads to isolate all cDNA molecules and 0.4X to enrich for higher molecular weight. DNA concentration was accurately measured using Quant-iT Picogreen (Thermo Fisher) and each sample was pooled in a mass ratio of 4:1 favoring high molecular weight cDNA. All barcoded samples were pooled in nearly equal amounts to produce a single library prepared using the SMRTbell Template Prep Kit 1.0-SPv3 (Pacific Biosciences, Menlo Park, CA) according to the manufacturer’s protocol. The library was supplied to the University of Washington PacBio sequencing facility and analyzed using a PacBio Sequel system with a SMRT Cell 8M recorded for 30 hours. Raw movies were decoded to full length non-concatenated cDNA sequences with circular consensus averaging using the IsoSeq3 software pipeline. Data were deposited to the NCBI Sequence Read Archive (BioProject ID PRJNA785352).

### Bioinformatic analysis of transcriptomic data

Pacbio Iso-Seq reads were greedily clustered using isONclust (Sahlin and Medvedev, 2020) and transcript abundance estimated as the read count per cluster and normalized to the total number of reads per sample, reported as transcripts per million (TPM). For each read, all possible coding sequences from an initiator Met to stop codon that coded for at least 30 amino acids were identified from the three forward reading frames (Iso-Seq reads are stranded), and then filtered to retain only the set of longest possible non-overlapping coding sequences per read. Potential coding sequences and their protein translations were aggregated from all reads. Protein translations supported by at least 2 reads were included in a search database for proteomic analysis. This putative protein data was also analyzed by protein BLAST to Uniref90 (accessed Aug 28, 2020) (Camacho et al., 2009) and leader signal peptide sequences identified by signalp (Armenteros et al., 2019)

### Proteomics sample preparation and liquid chromatography mass spectrometry

Plethodontid spermatophores stored in RNAlater were centrifuged at 10,000 x *g* for 10 min, RNAlater removed, the pellet resuspended in 50 μL 10% (w/v) SDS/50 mM TEAB/20mM DTT, and incubated at 65°C for 30 min. The samples were briefly chilled on ice and disulfide bonds alkylated by addition of 4.6 μL 0.5M iodoacetamide with incubation in the dark at room temperature for 30 min. Samples were acidified by addition of 5.6 μL 12% phosphoric acid, mixed with 0.35 mL 90% methanol/100mM TEAB before the entire solution was applied to a suspension trap (S-Traps; Protifi LLC) by centrifugation at 4,000 x *g* for 30 sec, and three washes of 0.4 mL 90% methanol/100mM TEAB were applied using the same centrifugation conditions. S-Traps were loaded with 125 μL trypsin solution (50 ng/μL in 50mM TEAB; Promega) and incubated at 47°C for 2 hours. Peptides were extracted from S-Traps by serial application of 50 mM TEAB, 0.2% formic acid, and 50% acetonitrile/0.2% formic acid in 80 μL volumes followed by centrifugation at 1,000 *x g* for 1 min. Resulting peptides were dried with vacuum centrifugation and resuspended in 0.1% formic acid at concentrations of ~1 μg per 3 μL immediately prior to mass spectrometry acquisition. Four biological peptide samples from each species were pooled for mass spectrometry analysis.

Tryptic peptides were separated with a Thermo Easy nLC 1200 and emitted into a Thermo Exploris 480 using a 75 μm inner diameter fused silica capillary with an in-house pulled tip. The column was packed with 3 μm ReproSil-Pur C18 beads (Dr. Maisch) to 28 cm. A Kasil fritted trap column was created from 150 μm inner diameter capillary packed to 1.5 cm with the same C18 beads. Peptide separation was performed over a 90-minute linear gradient using 250 nL/min flow with solvent A as 0.1% formic acid in water and solvent B as 0.1% formic acid in 80% acetonitrile. For each injection, 3 μL (400-900 m/z injections) or 5 μL (900-1,000 m/z injections) was loaded. Following the approach described in (Pino et al., 2020), six gas-phase fractionated data independent acquisition (GPF-DIA) experiments were acquired of each sample (120,000 precursor resolution, 30,000 fragment resolution, fragment AGC target of 1000%, max IIT of 55 ms, NCE of 27, +3H assumed charge state) using 4 m/z precursor isolation windows in a staggered window pattern with optimized window placements (i.e., 398.4 to 502.5 m/z, 498.5 to 602.5 m/z, 598.5 to 702.6 m/z, 698.6 to 802.6 m/z, 798.6 to 902.7 m/z, and 898.7 to 1002.7 m/z).

### Mass spectrometry data processing

Putative proteins sequenced by at least 2 reads in the combined plethodontid gonad transcriptome (300,472 total sequences) were digested *in silico* to create all possible +2H and +3H peptides between 7 and 30 amino acids with precursor m/z within 396.43 and 1,002.70, assuming up to one missed tryptic cleavage. Peptide fragmentation and retention time predictions for these peptides were made with the Prosit webserver (2020 HCD model, 2019 iRT model) (Gessulat et al., 2019) and collected in a spectrum library using the approach presented in Searle et al. (Searle et al., 2020). DIA data was demultiplexed (Amodei et al., 2019) with 10 ppm accuracy after peak picking in ProteoWizard (version 3.0.18113) (Chambers et al., 2012). Library searches were performed using EncyclopeDIA (version 1.2.2) (Searle et al., 2018), which was set to search with 10 ppm precursor, fragment, and library tolerances, considering both B and Y ions and assuming trypsin digestion. Detected peptides were filtered to a 1% peptide and protein-level false discovery rate. Mass spectrometry data have been deposited on the ProteomeXchange Consortium via the PRIDE (Perez-Riverol et al., 2019) partner repository with the dataset identifier PXD030143. Peptides were organized into protein groups based on parsimonious analysis of sequence clusters (i.e. “genes”) from the gonad transcriptome, with multiple clusters consolidated into an individual protein group if at least two identified peptides were shared between clusters. If at least three peptides were detected for a protein group, protein abundance was estimated as the mean ion intensity of the three most intense peptides (Silva et al., 2006).

### Molecular evolutionary analysis of courtship pheromones and sperm paralogs

Homologs of pheromone proteins were identified in the gonad proteomics database by protein BLAST using sequences of PRF, PMF, and SPF for which there is proteomic evidence in plethodontid mental glands (Wilburn et al., 2012;Wilburn et al., 2014b;Doty et al., 2016). Given the high sequence divergence between TFP homolog families, additional TFP-like sequences were identified in the gonad proteomics database by searching with the regular expression “.{2}C.{5,30}C.{2,20}C.{5,30}C.{2,20}C.{5,30}CC.{4}CN” using a custom Python script. PMF and all TFP-like sequences in the sperm proteomes were compared to other TFP families using sequences from representative vertebrate species if the TFP family was amphibian-specific (Amplexin, Prod1), detected by protein BLAST with the sperm paralog of PMF regardless of e-value CD59, Ly6E, SLURP1, Prostate Stem Cell Antigen) or is associated with fertilization (SPACA4, Bouncer). Homologs of SPF were similarly curated based on homologs identified to protein BLAST to sequences of representative vertebrate species with a bias towards amphibian SPF-like sequences from the references (Van Bocxlaer et al., 2015;Maex et al., 2016;Bossuyt et al., 2019). PRF phylogenetics focused only on sequences identified in the gonad proteomics database. Protein groups were aligned using MAFFT (v7.475) with E-INS-I (Katoh and Standley, 2013), and maximum likelihood trees were estimated using raxml-ng (v.0.9.0) (Kozlov et al., 2019) with 100 starting trees (50 random, 50 parsimony) using the LG amino acid substitution matrix and gamma distributed rate heterogeneity. Branch support was computed as the transfer bootstrap effect (TBE) (Lemoine et al., 2018) from 200 bootstrap trees.

## Results

### Transcriptomic analysis of plethodontid testis and ovary

Transcriptomic analysis of RNA isolated from testis and ovary of *P. shermani* and *D. ocoee* was performed using PacBio Iso-Seq to enumeate potential proteins within plethodontid gametes. While short-read sequencing technologies require that reads be assembled (either *de novo* or by genomic alignment) to reconstruct protein coding sequences, PacBio Iso-Seq utilizes long read single molecule sequencing with circular adapters and consensus averaging to produce high-quality, full-length transcript sequences without assembly (Gonzalez-Garay, 2016). Using RNA isolated from testis and ovary samples of both species in biological duplicate (n = 8), molecularly barcoded cDNA was synthesized, pooled, and sequenced using a single PacBio SMRT cell to generate a total of ~3 million reads with similar proportions for each sample (Table 1). Reads were clustered based on sequence similarity by the greedy algorithm isONclust into putative gene groups, and for simplicity we will refer to each read cluster as a gene. In both *P. shermani* and *D. ocoee*, comparison of relative gene expression between testis and ovary revealed dramatically different expression patterns between the gonads. Both testis and ovary express several thousand sex-biased genes (defined as >10-fold difference between the tissues), but gene expression patterns in testis are more extreme by multiple metrics. First, the mostly highly expressed genes in testis are found at ~5-10X higher levels compared to the most abundant transcripts in ovary. Second, highly abundant transcripts in testis are almost always sex-biased, while this is less common in ovary (Figure 1, Table 2). Third, testis expresses many more thousands of genes compared to ovary, most of which are at very low abundance (i.e. singleton reads) (Figure S1). This observation is consistent with genome wide transcription within testis as is common in other vertebrates and may enhance genomic fidelity through increased proofreading (Xia et al., 2020). Fourth, both within and between species, there is greater variability in the expression profiles of testis samples compared to ovary (Figure S2). Such transcriptional variation in testis may relate to the percent of testicular RNA derived from sperm versus other cell types, as well as the proportion of sperm at different stages of spermatogenesis. Both variables are known to seasonally vary from May through September (Woodley, 1994), with all samples in this study collected in August.

**Table 1.** Summary statistics of plethodontid gonad transcriptome

| Property | Quantity |
| --- | --- |
| Total number of reads | 3,096,431 |
| *P. shermani* testis reads | 719,087 |
| *D. ocoee* testis reads | 722,082 |
| *P. shermani* ovary reads | 831,814 |
| *D. ocoee* ovary reads | 823,448 |
| Mean length of full-length, non-concatamer reads | 2,321 |
| Number of sequence clusters | 166,909 |
| Mean number of reads per cluster | 18.6 |
| Number of non-singleton clusters | 60,007 |
| Mean number of reads per non-singleton cluster | 49.8 |
| Number of clusters with testis-biased expression (>10-fold) in *P. shermani* | 3,152 |
| Number of clusters with ovary-biased expression (>10-fold) in *P. shermani* | 2,405 |
| Number of clusters with testis-biased expression (>10-fold) in *D. ocoee* | 3,083 |
| Number of clusters with ovary-biased expression (>10-fold) in *D. ocoee* | 3,298 |

**Table 2.**
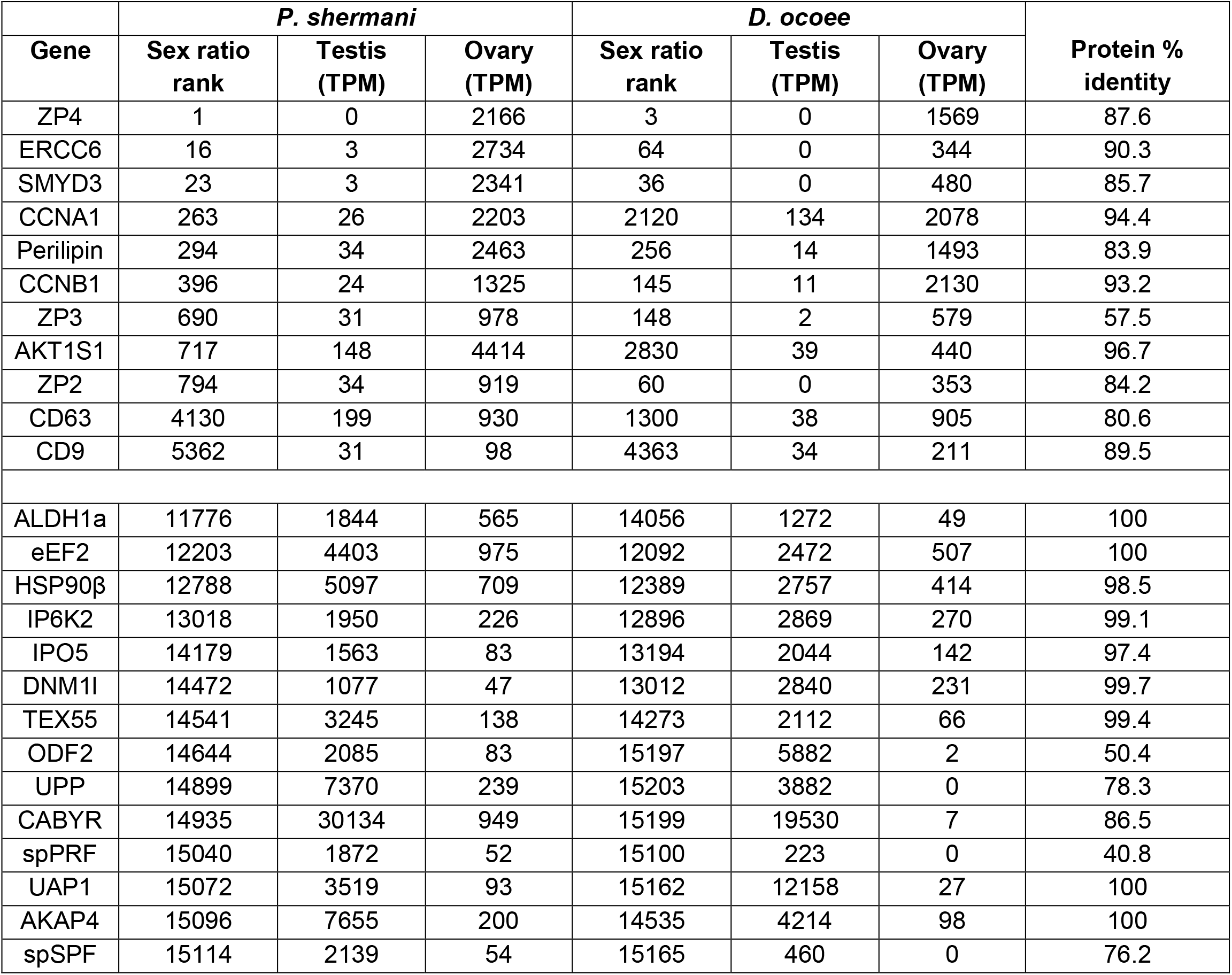
Transcriptome abundance of major sex-biased genes

| Gene | *P. shermani* |  |  | *D. ocoee* |  |  | Protein % identity |
| --- | --- | --- | --- | --- | --- | --- | --- |
|  | Sex ratio rank | Testis (TPM) | Ovary (TPM) | Sex ratio rank | Testis (TPM) | Ovary (TPM) |  |
| ZP4 | 1 | 0 | 2166 | 3 | 0 | 1569 | 87.6 |
| ERCC6 | 16 | 3 | 2734 | 64 | 0 | 344 | 90.3 |
| SMYD3 | 23 | 3 | 2341 | 36 | 0 | 480 | 85.7 |
| CCNA1 | 263 | 26 | 2203 | 2120 | 134 | 2078 | 94.4 |
| Perilipin | 294 | 34 | 2463 | 256 | 14 | 1493 | 83.9 |
| CCNB1 | 396 | 24 | 1325 | 145 | 11 | 2130 | 93.2 |
| ZP3 | 690 | 31 | 978 | 148 | 2 | 579 | 57.5 |
| AKT1S1 | 717 | 148 | 4414 | 2830 | 39 | 440 | 96.7 |
| ZP2 | 794 | 34 | 919 | 60 | 0 | 353 | 84.2 |
| CD63 | 4130 | 199 | 930 | 1300 | 38 | 905 | 80.6 |
| CD9 | 5362 | 31 | 98 | 4363 | 34 | 211 | 89.5 |
| ALDH1a | 11776 | 1844 | 565 | 14056 | 1272 | 49 | 100 |
| eEF2 | 12203 | 4403 | 975 | 12092 | 2472 | 507 | 100 |
| HSP90β | 12788 | 5097 | 709 | 12389 | 2757 | 414 | 98.5 |
| IP6K2 | 13018 | 1950 | 226 | 12896 | 2869 | 270 | 99.1 |
| IPO5 | 14179 | 1563 | 83 | 13194 | 2044 | 142 | 97.4 |
| DNM1I | 14472 | 1077 | 47 | 13012 | 2840 | 231 | 99.7 |
| TEX55 | 14541 | 3245 | 138 | 14273 | 2112 | 66 | 99.4 |
| ODF2 | 14644 | 2085 | 83 | 15197 | 5882 | 2 | 50.4 |
| UPP | 14899 | 7370 | 239 | 15203 | 3882 | 0 | 78.3 |
| CABYR | 14935 | 30134 | 949 | 15199 | 19530 | 7 | 86.5 |
| spPRF | 15040 | 1872 | 52 | 15100 | 223 | 0 | 40.8 |
| UAP1 | 15072 | 3519 | 93 | 15162 | 12158 | 27 | 100 |
| AKAP4 | 15096 | 7655 | 200 | 14535 | 4214 | 98 | 100 |
| spSPF | 15114 | 2139 | 54 | 15165 | 460 | 0 | 76.2 |

**Figure 1.**
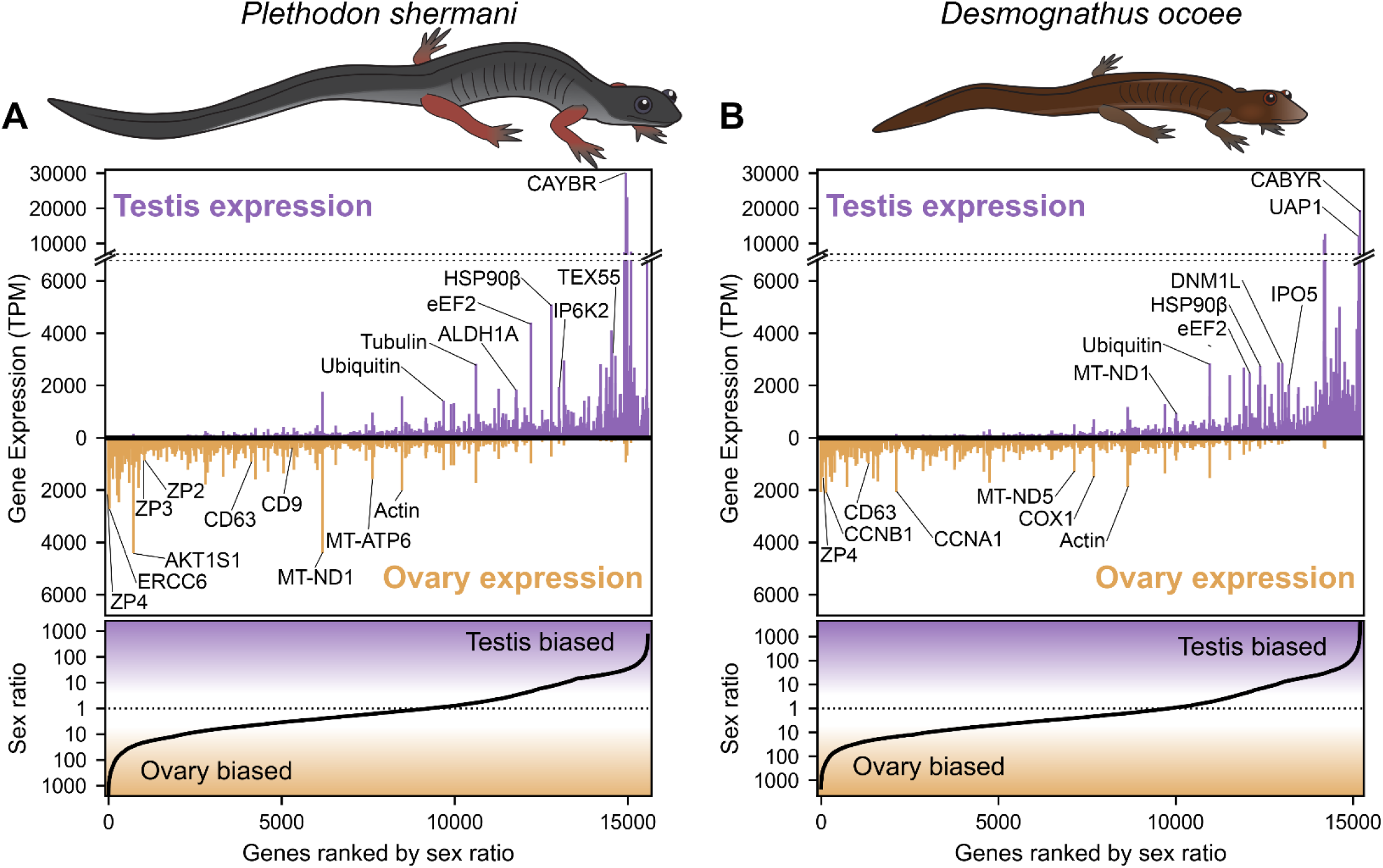
Abundance and sex bias of gene expression in plethodontid gonads. Comparison of the relative gene expression (measured as transcripts per million, TPM) between testis and ovary for genes identified in the plethodontid gonad transcriptome for (A) *P. shermani* and (B) *D. ocoee*. For each species, genes are ranked in order from most ovary-biased to most testis-biased with genes of interest annotated in the upper panel.

Examination of ovary-biased genes identified a few major functional categories common to vertebrate oocytes. The most evident example was zona pellucida (ZP) associated proteins: a family of secreted glycoproteins common to all animal oocytes that polymerize to form an extracellular egg coat that restricts sperm access to the egg plasma membrane. This egg coat is termed the zona pellucida (ZP) in mammals and the vitelline membrane (VM) in amphibians (Wassarman, 1999;Wilburn and Swanson, 2018) In both plethodontid species, homologs of the ZP genes *zp4, zp3*, and *zp2* were identified in the same gene expression rank order, with *zp4* being the first and third most ovary-biased genes in *P. shermani* and *D. ocoee*, respectively. This expression pattern more closely resembles the relative ZP protein abundances in the VM of *Xenopus spp*., where there is similar stoichiometry of ZP3 and ZP4 with lower levels of ZP2 (Lindsay et al., 2002) compared to the mammalian ZP where ZP3 and ZP2 are the major components with substantially less ZP4 and/or its close paralog ZP1. No other ZP genes were identified in the plethodontid transcriptome, including *zpax* and *zpd* that are minor components of the *Xenopus* VM, indicating that the plethodontid VM (and possibly the VM of other salamanders) may be similar but less complex than its anuran counterparts. Comparison of the *P. shermani* and *D. ocoee* sequences suggest that *zp3* may be evolving more rapidly with only 57.5% shared protein identity between the species, compared to ~85% identity for both *zp4* and *zp2*. Another interesting set of ovary biased genes were *cd9* and *cd63* that code for tetraspanin proteins associated with vesicular and cell fusion. In mammals, CD9 is highly expressed by oocytes and is considered a major candidate that facilitates sperm-egg fusion, while CD63 is more associated with exosomes found in follicular fluid produced by granulosa cells that surround oocytes and serve a different role in oogenesis (Jankovičová et al., 2015). The increased expression of *cd63* relative to *cd9* in the ovaries of both *P. shermani* and *D. ocoee* may support that salamander oocytes express both proteins, consistent with plethodontid ovaries having few if any supporting cells around developing eggs (D.B. Wilburn and P.W. Feldhoff, personal observations). Additional highly expressed ovary-biased genes common to vertebrate oocytes include *perilipin* which regulates lipid acquisition and lipid droplet formation during mammalian oogenesis (Yang et al., 2010;Zhang et al., 2014), and *smyd3* which codes for a histone methyltransferase which regulates expression of transcription factors critical for mammalian oocyte maturation and early embryonic development (Bai et al., 2016). Furthermore, one of the most abundant transcripts in *P. shermani* ovaries was a homolog of *ercc6* which in humans codes for a DNA binding protein involved in transcription-coupled DNA damage repair, and may play a role in preserving genomic fidelity in the salamander germline (Troelstra et al., 1992).

### Proteomic analysis of mature plethodontid sperm

Following meiosis and compaction of the nuclear genome, vertebrate sperm and egg are usually transcriptionally silent. These cells differentiate via a complex cellular program that selectively translates transcripts using RNA binding proteins and waves of cytoplasmic polyadenylation (Belloc et al., 2008;Brook et al., 2009). Consequently, transcriptomic profiling of gonads primarily samples RNA from immature gametes (as well as other cell types in the tissue) that may not well represent their mature, fully differentiated forms. During plethodontid courtship, sperm is transferred by the male depositing a spermatophore on the substrate which the female will accept via her cloaca. By mating salamanders in the laboratory but interrupting courtship between spermatophore deposition and insemination, we exploited this system of external sperm transfer to collect spermatophores from *P. shermani* and *D. ocoee* and performed proteomic analysis of mature sperm. For each species, tryptic peptides were prepared from four spermatophores (each from a separate male salamander) and pooled in approximately equimolar amounts (based on total ion current) for proteomic analysis by mass spectrometry. Using the gonad transcriptome supplemented with mental gland cDNA sequences as a search database, we identified a total of 7287 peptides for 1648 protein groups in *P. shermani* sperm and 5308 peptides for 1157 protein groups in *D. ocoee* sperm. Protein groups, which are defined based on shared peptides identified in the sperm proteome (see methods), typically contain multiple genes from the transcriptome analysis as sequence variation in either the coding or noncoding regions may have led to the formation of multiple read clusters that contain similar predicted proteins.

For protein groups with at least 3 detected peptides, protein abundance was estimated using the mean ion intensity of the 3 most intense ions (Silva et al., 2006) (Figure 2). In both species, the most abundant protein was derived from genes with low transcript abundance and no significant BLAST hits. These small (~6.5 kDa), highly abundant proteins with ~50% of residues being arginine most likely function similarly to sperm protamines which replace nuclear histones following meiosis. The plethodontid protamine-like sequences are ~2-3X smaller than protamines of either mammals or *Xenopus spp*., and are devoid of cysteine residues that form intermolecular disulfide bonds and stabilize the mammalian sperm genome (Hutchison et al., 2017). Present at high levels in both the testis transcriptome and sperm proteome of both species were homologs of enzymes associated with uracil metabolism: uridine phosphorylase (UPP) and UDP-N-acetylhexosamine pyrophosphorylase (UAP1). UPP can reversibly interconvert uracil and ribose-1-phosphate into uridine and phosphate, while UAP1 converts uracil triphosphate (UTP) into UDP-N-acetylglucosamine or UDP-N-acetylgalactosamine that are substrates of protein glycosylation. In studies of human prostate cancer cell lines (Itkonen et al., 2013;Itkonen et al., 2015), activation of the androgen receptor (AR) increases expression of UAP1 such that similar regulatory machinery may be driving its high expression in plethodontid sperm. Also found at high levels in both the testis transcriptome and sperm proteome of both species were major proteins associated with the sperm flagellum, including multiple homologs of ODF2 that form outer dense fibers surrounding the flagellar axoneme (Zhao et al., 2018), which are further surrounded by a fibrous sheath largely comprised of AKAP4 and CABYR (Naaby-Hansen et al., 2002;Li et al., 2010). Notably, homologs of major mammalian sperm proteins associated with oocyte ZP or sperm-egg plasma membrane fusion, such as acrosin, SPACA4, or Izumo1 (Bianchi et al., 2014;Hirose et al., 2020;Fujihara et al., 2021) were observed at very low abundance in the testis transcriptome. Acrosin is the only example of these proteins that was observed in the *P. shermani* sperm proteome as the 483^rd^ most abundant protein, and none were observed in *D. ocoee* sperm.

**Figure 2.**
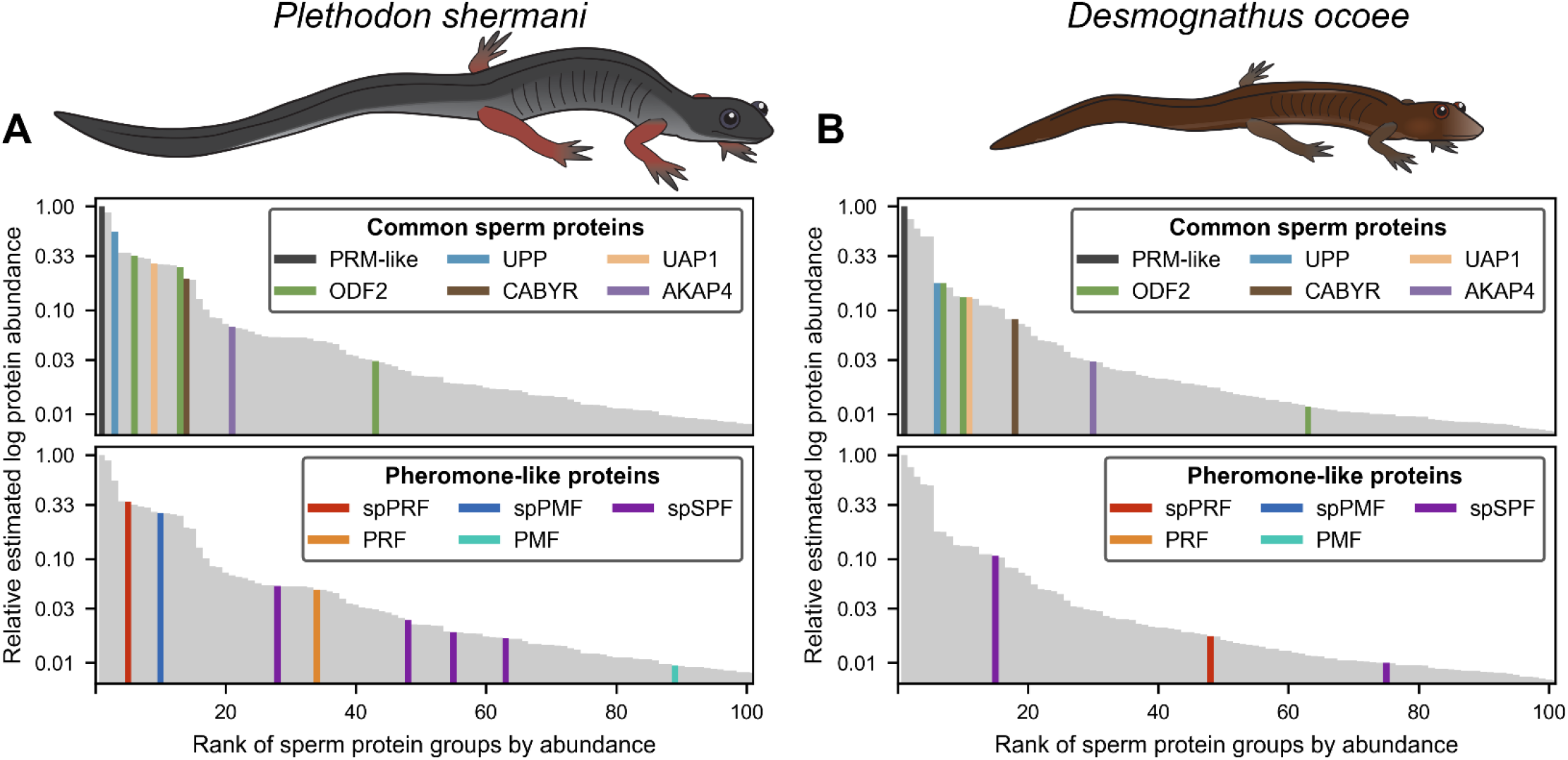
Relative abundance of major proteins in the plethodontid sperm proteome. Comparison of the estimated protein abundance (measured as the mean intensity of the three most intense peptides per protein group) for the top 100 proteins identified in the sperm of (A) *P. shermani* and (B) *D. ocoee*. For each species, estimated protein abundances were normalized to the most abundant identified protein (which was a novel protamine-like protein in both *P. shermani* and *D. ocoee*). Proteins are ranked according to their relative abundance. Common sperm proteins and pheromone-like proteins are separately highlighted relative to the protein abundance distribution for each species.

Beyond major structural and metabolic proteins, some of the most abundant proteins in the sperm proteomes of both *P. shermani* and *D. ocoee* were paralogs of the major mental gland pheromones SPF, PMF, and PRF. Based on signal peptide prediction, the sperm paralogs of PMF (spPMF) and SPF (spSPF) were estimated to be the most abundant secreted proteins in *P. shermani* and *D. ocoee*, respectively, while the sperm paralogs of PRF (spPRF) were surprisingly lacking signal peptides. The sperm expression patterns of these pheromone paralogs also mirror their mental gland expression within each species, with spPRF and spPMF being more abundant in *P. shermani* compared to spSPF which was more abundant in *D. ocoee*. Also surprising was the detection of the pheromone proteins themselves in the sperm proteome, but at much lower levels than their sperm-specific paralogs (*P. shermani*: [spPRF]/[PRF] ≈ 7, [spPMF]/[PMF] ≈ 30; *D. ocoee:* [spSPF]/[SPF] ≈ 27). Given the magnitude of these expression differences and known functions for the pheromones in regulating female courtship behavior (Rollmann et al., 1999;Houck et al., 2007;Wilburn et al., 2015), we hypothesize that pheromone expression in sperm is the result of leaky gene expression. As both pheromone transcript and protein levels are correlated with seasonal elevation in plasma androgen levels (Woodley, 1994;Wilburn et al., 2019), the sperm paralogs likely rely on similar regulatory machinery. High levels of *spPRF* and *spSPF* in testis of both *P. shermani* and *D. ocoee* was consistent with their elevated protein expression, while *spPMF* was much less abundant than expected with only 7 transcript reads acquired from all samples. This suggests that *spPMF* may be transcribed and/or translated at a different stage of sperm development relative to the other pheromone paralogs. Interestingly, no pheromone sequences were identified in the transcriptome.

### Molecular evolutionary analysis of plethodontid courtship pheromones and their sperm paralogs

To investigate the evolutionary dynamics between courtship pheromones and their sperm paralogs, detailed phylogenetic molecular evolutionary analyses were performed on each of the three pheromone families: PMF, SPF, and PRF. PMF is a family of small ~7 kDa proteins with homology to the TFP superfamily, a highly diverse family of vertebrate proteins whose functions include examples such as neuro- and cytotoxins within snake venoms (Fry, 2005), spatial signaling during amphibian limb regeneration (Garza-Garcia et al., 2009), membrane receptors in mammalian tissue reorganization (Blasi and Carmeliet, 2002), and regulation of the complement system (Davies et al., 1989). Despite this extraordinary array of functions, TFPs are defined by their namesake “three finger” topology that includes a 2- and 3-stranded β-sandwich stabilized by a core of 8 cysteine residues that adopt an invariant disulfide bonding pattern. Some TFPs include an extra pair of cysteines that form an extra disulfide bond within the first finger that does not affect the arrangement of the 8-cysteine core, and we will refer to these extra disulfide containing proteins as 10C-TFPs compared to 8C-TFPs. While both the lengths of the β-strand and the exact spacing of the first 5 (or 7) cysteines can vary between homologs, the only strictly conserved sequence motif in the TFP domain is the last 8 residues being CCXXXXCN, with the terminal asparagine participating in two critical H-bonds that stabilize the β-sandwich structure (Galat et al., 2008). While PMF is homologous to and possesses the sequence characteristics of 8C-TFPs, the solution structure for the most abundant PMF isoform in *P. shermani* revealed a novel protein topology with an altered disulfide bonding pattern (Wilburn et al., 2014a). This major PMF is only one out of more than 30 isoforms in the *P. shermani* mental gland that are highly polymorphic with only ~30% average identity between amino acid sequences. The PMF coding sequence evolves extremely rapidly while the flanking untranslated regions (UTRs) within PMF mRNAs are unusually highly conserved with ~98% nucleotide identity in both the 5’ and 3’ UTR sequences. Analysis of the PMF UTR sequences identified 3 paralogous classes of PMF genes (Class I, II, and III) that were likely present in the last common ancestor of all plethodontid salamanders (Wilburn et al., 2012). Most *P. shermani* PMFs are of Class I or III (~50% total pheromone by mass) while the *D. ocoee* mental gland expresses only a single Class I PMF at low levels (<5% total pheromone) (Chouinard et al., 2013;Doty et al., 2016).

The sequence of spPMF possesses several notable characteristics when compared to the major pheromone PMFs of *P. shermani*, *D. ocoee*, and an intermediate species *Plethodon cinereus* (Figure 3A). While most PMFs are highly negatively charged, spPMF has an estimated net charge of +14 at physiological pH with 20 out of the 71 residues being lysine and none are arginine. The density of these positive charges is especially high in finger 3, which also poses a potential N-glycosylation site following Cys 5. Despite the large number of lysine residues, spPMF lacks a lysine at position 30 that normally would form a salt bridge with E35 to stabilize finger 2. The C-terminus of spPMF also contains a unique three residue extension (TGK) past the conserved asparagine, with peptide coverage from the proteomics data confirming that these additional residues are present in the mature protein. There was no detectable sequence similarity between the spPMF UTRs to those of Class I, II, or III. Protein BLAST searches of spPMF returned PMF as the top hit but with only modest confidence (e-value ~ 1e-5), and given the extreme sequence differences, it was unclear if spPMF might be more closely related to another 8C-TFP family. Phylogenetic analysis was performed with a diverse panel of TFP sequences that included all characterized PMFs, all TFP sequences in the sperm proteomes (including spPMF), as well as representatives of other amphibian-specific TFPs (Amplexin, Prod1), families detected by BLAST searches of plethodontid gonad-associated TFPs regardless of e-value (CD59, Ly6E, SLURP1, Prostate Stem Cell Antigen), and TFPs with known reproductive functions (SPACA4, Bouncer). Based on the estimated maximum likelihood tree, spPMF was found on a relatively long branch adjacent to the pheromone PMFs, supporting that they share a more recent comparison ancestor compared to all other analyzed TFPs (Figure 3B). Since the gene duplications that gave rise to the three PMF classes likely occurred in the last common ancestor of plethodontid salamanders, this topology suggests that the duplication event separating PMF and spPMF must be similarly ancient. Sister to PMF/spPMF are the other amphibian specific families Prod1 and Amplexin-like proteins, both of which are present in plethodontid salamanders such that a single gene duplication of an ancestral amphibian TFP may have given rise to the ancestor of PMF and spPMF. As presence of a mental gland are considered the ancestral condition of plethodontid salamanders (Sever et al., 2016), it is unclear in which tissue such a PMF-like ancestor may have originated but likely evolved to function as both a pheromone and a sperm protein at similar evolutionary points.

**Figure 3.**
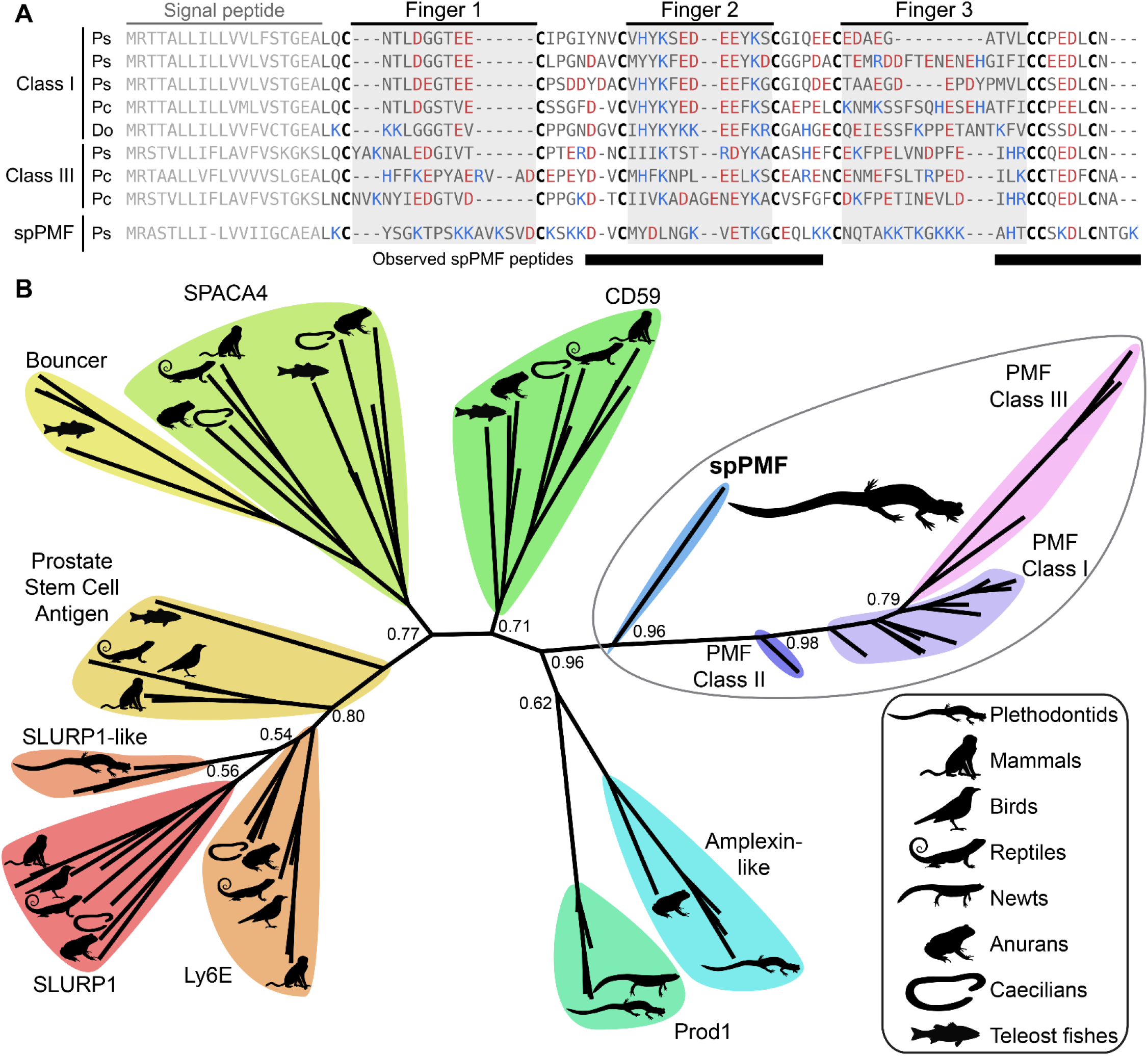
Comparison of sperm PMF (spPMF) to pheromone PMF and other TFP proteins. (A) Alignment of spPMF with major class I and III PMF proteins found in the mental gland proteomes of *P. shermani* (Ps), *P. cinereus* (Pc), and *D. ocoee* (Do). Bars beneath the alignment denote sequence coverage of tryptic peptides identified in the *P. shermani* sperm proteome. Cysteine residues that define the relative spacing of the three fingers in TFPs are denoted in bold, with basic and acidic residues colored in blue and red, respectively. The signal peptide that is not part of the mature protein sequence is greyed out. (B) Maximum likelihood tree of spPMF and related TFP proteins in vertebrates, with branch support reported for major nodes between TFP paralogs. The three classes of PMF pheromones form a single, plethodontid-specific clade with spPMF, suggesting that a common ancestral gene gave rise to both the mental gland and sperm proteins independent of other major TFP families.

SPF is an incredibly ancient pheromone family with homologs that influence female mating behavior in both caudates (salamanders and newts) and anurans (frogs and toads), implying that SPF has played a critical role in amphibian reproduction for ~360 million years (Bossuyt et al., 2019). An ancient gene duplication of SPF produced two families, SPFα and SPFβ, that were both present in the last common ancestor of amphibians, with subsequent duplications giving rise to multiple paralogs within all examined taxa (Janssenswillen et al., 2014). The courtship pheromones of plethodontid salamanders are derived from SPFα while most characterized pheromones in other caudates and frogs are descended from SPFβ (Maex et al., 2016). Like PMF, SPF is also homologous to the TFP superfamily except that it is part of a distantly related subfamily that arose from a tandem duplication of 10C-TFPs such that it has two TFP-like domains. In the major SPF of *D. ocoee*, both TFP-like domains have highly altered disulfide bonding patterns that likely produces an altered topology and are not classical TFPs (Doty et al., 2016). To characterize the relationship between SPF and spSPF, a maximum likelihood tree was generated based on SPF homologs identified by BLAST for all major vertebrate clades where SPFs from salamanders and newts were overrepresented to improve resolution around plethodontid sequences (Figure 4A). Manual examination of these sequences identified five variations of cysteine spacings that ranged in complexity from two stereotypical 10C-TFP patterns to the highly altered arrangement determined for the major *D. ocoee* SPF (Figure 4B). Teleost fish exclusively had predicted proteins with two canonical 10C-TFP patterns – the likely ancestral condition of these SPF-like proteins – and the maximum likelihood tree was rooted on the branch of fish homologs with true two domain TFPs (2D-TFPs). All spSPF sequences shared the same cysteine spacing where the first TFP-like domain had a normal 10C-TFP pattern and the second domain was missing the conserved C-terminal CCXXXXCN motif critical for stabilizing the β-sandwich structure. While the plethodontid pheromone SPFs formed a single clade nested among other SPFα sequences, spSPF sequences identified in the gonad transcriptome localized to five well supported clades (>90% bootstrap support): 3 in SPFβ and 2 in SPFα. The most abundant spSPF proteins in both *P. shermani* and *D. ocoee* sperm was localized to one of the SPFβ clades with close homology to SPF12 and SPF13 from the Mexican axolotl (*Ambystoma mexicanum*). Maex et al. (2016) identified multiple new SPF proteins in male axolotl salamanders with transcriptomic analysis suggesting that most isoforms were expressed by male cloacal glands. However, SPF12/13 were unusual compared to other isoforms with ~100-5000x lower transcript abundance in cloacal glands despite strong proteomic evidence of the proteins in water that was inhabited by male salamanders during mating. The close phylogenetic relationship of axolotl SPF12/13 with spSPF of both *P. shermani* and *D. ocoee* suggests that these axolotl proteins may instead be sperm proteins rather than pheromones, and would place the origin spSPF at the base of modern salamanders (~160 million years ago) (Shen et al., 2015). In contrast to PMF where a single plethodontid gene likely gave rise to both the pheromone and sperm paralogs, the role of SPF in these two reproductive contexts arose independently from ancient paralogs that appears to have duplicated ~360 million years ago in the last common ancestor of amphibians.

**Figure 4.**
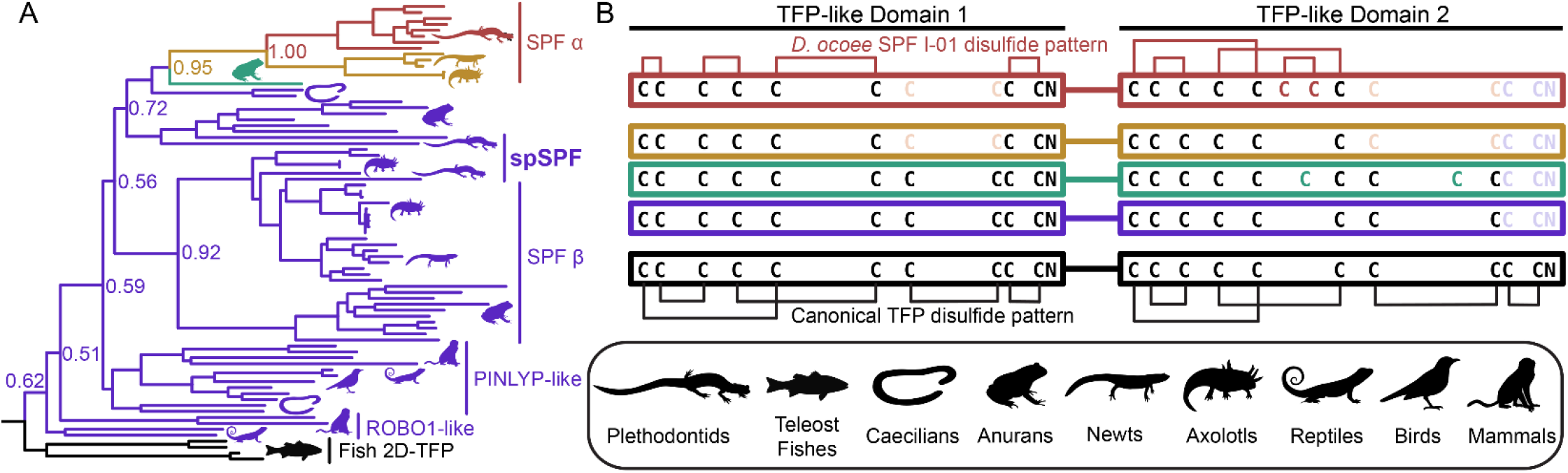
Evolution of SPF protein evolution relative to 2D-TFPs. A maximum likelihood tree of SPF-like protein sequences from representative vertebrates with overrepresentation of salamanders and newts, where numbers denote branch support of major nodes (A). Tree branches are colored according to the arrangement of cysteine residues in the underlying sequence alignment as denoted in (B). Black denotes the ancestral condition of two tandem 10C-TFP sequences that likely adopt the canonical disulfide pattern, and additional colors denoting sequential cysteine loss (light color text) and gain (dark color text), with the empirically determined disulfide pattern of *D. ocoee* SPF-I01 shown in red (Doty et al., 2016). The three is rooted on the branch of teleost fish proteins that are the only available sequences with the ancestral 2D-TFP cysteine spacing. Plethodontid pheromone SPF sequences are exclusively found within the red SPFα clade, with different spSPF paralogs being found in both the upper SPFα and lower SPFβ clades that share the purple cysteine pattern. The phylogenetic separation of the plethodontid pheromones from spSPF supports that each protein family was independently derived from gene duplications present in the last common ancestor of amphibians ~360 million years ago.

While SPF and PMF are ancestral to all plethodontid salamanders (~66 million years ago), PRF is a relatively young pheromone family related to α-helical cytokines and only present in the mental gland of *Plethodon spp*. from eastern North America that evolved ~17 million years ago (Shen et al., 2015). Detection of spPRF in both the transcriptome and proteome of *D. ocoee* (separated from *P. shermani* by ~43 million years) implies that spPRF is much older than the pheromone paralogs and likely originated as a sperm protein. BLAST searches of both spPRF and PRF to annotated vertebrate genomes identified cardiotrophins as the most similar homologs by sequence. Cardiotrophin-like sequences were identified with low but testis-biased expression in the gonad transcriptomes of both *P. shermani* and *D. ocoee*, supporting the hypothesis that the common ancestor of these proteins may have also had male-biased expression that facilitated the co-option of spPRF from cardiotrophins for sperm function. A maximum likelihood tree was estimated from all cardiotrophin and spPRF sequences identified in the gonad transcriptome in addition to representative PRF pheromone sequences from each major clade of eastern *Plethodon spp*. The cardiotrophin-like sequences from *P. shermani* and *D. ocoee* testis were found on a single long branch that was used to root the phylogeny (Figure 5A). The first branch in the PRF/spPRF clade separated all *Plethodon* sequences from *D. ocoee* spPRF, consistent with a sperm origin for PRF-like proteins. Within *Plethodon*, spPRF sequences were found in three separate clades and named according to homology with pheromone PRF types. Prior molecular evolutionary analysis identified two families of pheromone PRF: PRF-A which is common to all eastern *Plethodon spp*. and PRF-B which is only found in species closely related to *P. cinereus* with transdermal pheromone delivery (Wilburn et al., 2014b). As expression of PRF-B was only found in species with the more ancestral delivery system, it was hypothesized to be older than PRF-A (Watts et al., 2004). Instead, the updated phylogeny with spPRF sequences strongly supports that PRF-A and PRF-B arose independently from different spPRF paralogs termed spPRF-A and spPRF-B, respectively (Figure 5A). We were unable to distinguish the relative abundance of spPRF-A and spPRF-B in the *P. shermani* sperm proteome as most of the high intensity spPRF peptides are common to both paralogs. The third clade of spPRF termed spPRF-C is likely ancestral to spPRF-A/spPRF-B and is likely at much lower abundance, as few reads were identified in the testis transcriptome and no peptides unique to spPRF-C were detected in the sperm proteome. Compared to the unique histories of SPF/spSPF evolving independently and PMF/spPMF likely sharing a single origin, a third pattern is observed for PRF/spPRF where a family of sperm proteins was independently co-opted multiple times for pheromone activity.

**Figure 5.**
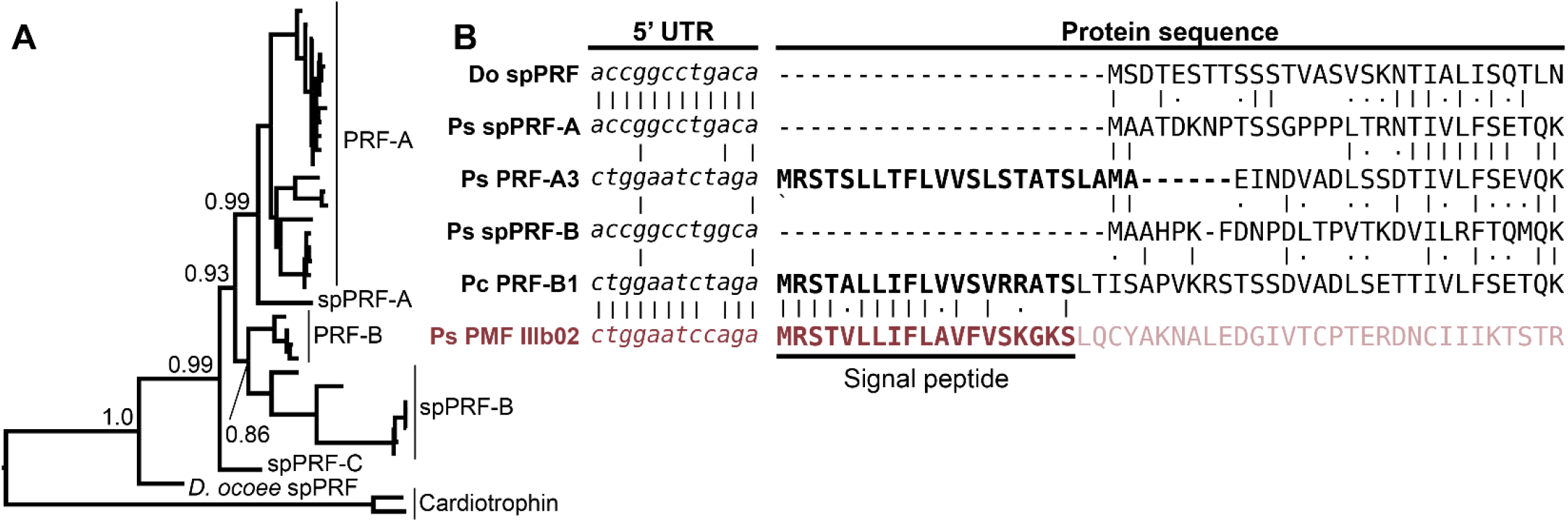
Multiple independent origins of pheromone PRF and recombination with PMF. (A) A maximum likelihood-based phylogeny of PRF and spPRF with numbers denoting branch support of major nodes and rooting based on cardiotrophin-like sequences from *P. shermani* and *D. ocoee*. The topology supports that an ancestral spPRF gene repeatedly duplicated into at least three families in *Plethodon*, and subsequent gene duplications of spPRF-A and B independently gave rise to PRF-A and B pheromone families. (B) Alignment of select PRF and PMF sequences around the translation start site of each protein. Despite high similarity in 5’ UTR and N-terminal protein sequence of spPRF from *D. ocoee* (Do) and *P. shermani* (Ps), representative PRF-A and B sequences from *P. shermani* and *P. cinereus* (Pc), respectively, have a distinct 5’ UTR sequence in addition to a signal peptide. This new PRF leader sequence is highly similar to that of PMF class III, such that it was likely acquired via recombination following the independent duplications of spPRF-A and B that gave rise to PRF-A and B, respectively.

In contrast to the pheromones and their sperm paralogs discussed so far, all spPRF sequences lack a signal peptide that would normally target it for secretion indicating that spPRF may be an intracellular protein. Peptides that include the initiator methionine of both spPRF-A and spPRF-B were observed in the *P. shermani* sperm proteome confirming that there is not a cryptic N-terminal signal peptide. To the best of our knowledge, spPRF is the first cytokine-like protein identified without a signal peptide. Presence of signal peptides in the testis-biased cardiotrophin-like sequences supports that spPRF lost its signal peptide following duplication from a cardiotrophin-like ancestor. While PMF and SPF require the oxidizing environment of the endoplasmic reticulum for proper disulfide bond formation, PRF adopts a predominantly α-helical fold without disulfides that could fold independently in cytoplasm (Houck et al., 2008). Secondary structure prediction of spPRF supports that it forms a similar α-helical topology and has no additional cysteine residues. Interestingly, while spPRF may be intracellular, both PRF-A and PRF-B have signal peptides despite independently arising from spPRFs lacking them. It was noticed that the PRF signal peptides were similar in amino acid sequence to those of PMF. Based on unpublished genome walking experiments, the gene structure of PMF includes 3 coding exons homologous to other TFP genes where ~90% of the signal peptide is coded by the first exon. Based on the shared sequence similarity and the high co-expression of PRF and PMF in *Plethodon* mental glands, we hypothesized that the PRF signal peptide may have been co-opted from a PMF gene. Manual alignment of the signal peptides between PRF-A/B and PMF classes I-III found the highest similarity between PRF-B and PMF Class III. Expansion of the alignment into the 5’ UTR immediately upstream revealed almost identical sequences between PRF-A, PRF-B, and PMF Class III that was distinct from spPRFs of both *P. shermani* and *D. ocoee* (Figure 5B). As the UTRs are ~98% identical between PMF paralogs of the same class, this high level of conservation in the PRF and PMF class III 5’ UTR strongly supports that an ancestral spPRF gene duplicated and recombined with a class III-like PMF to ultimately become PRF expressed by the mental gland. Given the independent origins of PRF-A and PRF-B, and the sequences of their 5’ UTR and signal peptides relative to PMF class III, our hypothesis as to the order of events is that spPRF-B duplicated and recombined with PMF class III to yield PRF-B, followed by spPRF-A duplicating and recombining with PRF-B to produce PRF-A.

## Discussion

Gametic fusion is an essential biological process for reproduction of most animal species, yet despite more than a century of research, it largely remains a molecular enigma as to *how* these highly specialized cells mechanistically accomplish this feat with species-specificity. The earliest studies of fertilization were in external fertilizing marine invertebrates where large numbers of gametes could be easily isolated for biochemical fractionation, and research continues for classic models such as sea urchins and abalone where the molecular interactions of sperm and egg proteins are best characterized (Wilburn et al., 2019;Wessel et al., 2021). Advances in gene editing technologies have elevated genetic knockouts as one of the primary tools to identify gamete recognition proteins, especially in mammals, but high rates of gene duplication and functional redundancy in gametic proteins can confound such studies (Baba et al., 1994;Isotani et al., 2017;Hirose et al., 2020). Due to these many challenges, there are currently only 4 *bona fide* pairs of interacting sperm-egg proteins identified in any animal species (Wilburn and Swanson, 2016). The extraordinarily fast evolution of reproductive proteins also facilitates the birth and death of new fertilization genes, resulting in gametic proteins often being shared by only closely related taxa and limiting the inferences that can be made about more distantly related species. As such, there is a need within reproductive biology to expand the taxonomic breadth of model systems and for more attention paid to clades that have been historically understudied. As a prime example of this myopia, amphibians are a major class of vertebrates with more than 6,000 species that have evolved over ~360 million years, and yet molecular characterization of amphibian gametes has been limited to a single genus of frogs (*Xenopus spp*.). In this study, we have provided the first detailed transcriptomic and proteomic analysis of gametes for one of the other major amphibian types: salamanders.

One of the most unusual qualities of plethodontid salamanders is the enormous size of their genomes: compared to the ~3 billion base pair haploid genome of humans, the sizes of the *P. shermani* and *D. ocoee* haploid genomes are ~27 billion and ~17 billion base pairs, respectively (Herrick and Sclavi, 2014). Plethodontid genomes contain high levels of long retrotransposons in intronic and intergenic regions (Sun et al., 2012), and this enormous genome likely creates unique challenges during gametogenesis such as proper meiotic chromosome segregation, maintenance of genome fidelity, as well as regulation of transcription and splicing of sex-specific genes with very large introns. Two of the most abundant ovary-biased transcripts code for the DNA repair protein ERCC6 and the histone methyltransferase SMYD3 which aid in these processes. The most abundant in sperm protein of both species is an extremely positively charged ~6.5 kDa protein that resembles protamines of other vertebrates and likely functions in replacement of histones as scaffolding for the nuclear genome. These protamine-like proteins are smaller and more arginine rich than their likely mammalian and *Xenopus* analogs, making them likely intrinsically disordered absent DNA. DNA packaged with poly-Arg is more dense than when packaged with poly-Lys (DeRouchey et al., 2013), and there may be a high premium on more tightly condensing the nuclear genome in the relatively small sperm cell. A more efficient system of nuclear packaging may also more effectively inhibit transcription in developing sperm, and if so, this may help explain the discrepancy in transcript and protein abundance in testis versus mature sperm, respectively.

For both plethodontid species, most genes with ovary biased expression coded for structural and metabolic proteins common to animal oocytes and we found no clear examples of salamander-specific innovations. Among the structural proteins, we were particularly interested in ZP genes that code for proteins in the VM, the extracellular egg coat that functions as a molecular gate to restrict both the number and species of sperm that may fertilize an oocyte. The plethodontid VM is likely similar in composition to the *Xenopus* VM with ZP4 and ZP3 being the major glycoproteins with smaller amounts of ZP2. Compared to other vertebrate clades, the amphibian VM is most biochemically similar to the mammalian ZP, and mouse ZP proteins are even able to integrate into the *Xenopus laevis* VM when synthetic mRNAs are microinjected into developing oocytes (Doren et al., 1999). While the molecular mechanisms of both mammalian ZP and *Xenopus* VM dissolution are poorly characterized, a few sperm proteins that physically interact with the mammalian ZP have been identified (Mori et al., 1993), although we found no homologs of these proteins in either the gonad transcriptome or sperm proteomes. Recently, a membrane-bound 10C-TFP called SPACA4 was implicated in mammalian ZP dissolution when sperm from *spaca4*-null mice could only fertilize oocytes without a ZP (Fujihara et al., 2021). While the precise function of SPACA4 remains unknown, the possible role of a TFP in ZP dissolution is intriguing given that spPMF and spPRF are likely the most abundant secreted sperm proteins in *P. shermani* and *D. ocoee*. The extreme positive charge of spPMF is also comparable to highly positively charged fertilization proteins in the marine mollusk abalone that mediate both egg coat dissolution and egg-sperm plasma membrane fusion (Wilburn et al., 2019). SPACA4 was present in the gonad transcriptome at very low levels and no peptides were detected in the sperm proteome of either species.

The evolutionary dynamics of the plethodontid courtship pheromones and their sperm paralogs spotlights the tremendous diversity of reproductive functions possible from only two protein families that are repeatedly subjected to gene duplication and sexual selection (summarized in Figure 6). SPF is the oldest known family of protein pheromones that evolved ~360 million years ago, derived from the ancestral tandem duplication of a 10C-TFP into the 2D-TFPs. Many paralogs of SPF are common in amphibian secretory glands which may be delivered by an array of courtship behaviors. For example, male palmate newts with aquatic reproduction release SPF into the water column from their cloaca and use their tails to propel these pheromones posteriorly to create a concentration gradient for chemotaxis (Van Bocxlaer et al., 2015). This passive broadcast of SPF pheromones contrasts with private transfer of SPF proteins when a male *D. ocoee* abrades the dorsum to a single female with his premaxillary teeth where pheromones are presumably diffused into the female bloodstream rather than stimulate the olfactory system (Houck et al., 2007). Recently, an extremely similar system was discovered in *Plectrohyla* frogs where males scratch the female dorsum with their teeth delivering SPF expressed by glands in the upper lip (Schulte et al., 2021). Despite the similarity in delivery system, the SPF pheromones of plethodontid salamanders and *Plectrohyla* frogs are independently derived from the SPFα and SPFβ families, respectively, that arose by gene duplication in a common ancestor of amphibians. As the highly abundant spSPF proteins in plethodontid sperm are also part of the SPFβ family, the original duplication that gave rise to these two distinctive plethodontid proteins also occurred at the base of amphibian evolution ~360 million years ago. The major spSPF proteins are only ~76% identical between *P. shermani* and *D. ocoee* such that sexual selection from coevolving female oocyte receptors may be driving rapid evolution, although both functional studies and sequencing of spSPF in additional plethodontid species are necessary to test this hypothesis. The evolution of spSPF is one example of at least three independent recruitments of TFP-like proteins into vertebrate gametes. Another TFP recruitment of TFPs for reproduction is SPACA4 in tetrapod sperm and its close teleost fish homolog Bouncer which regulates species-specific egg-sperm fusion (Herberg et al., 2018). It is intriguing that closely related TFP proteins are expressed in both male and female gametes of different vertebrate clades but remains unknown if egg Bouncer represents an ancestral condition of these proteins or if Bouncer/SPACA4 are independent paralogs that happen to share a more recent common ancestor than other TFPs. The third TFP recruitment is PMF, a plethodontid-specific TFP-like family that seemingly evolved from an amphibian-specific TFP ancestor as both a sperm protein and courtship pheromone ~43 million years ago (Shen et al., 2015). Following the innovation of olfactory pheromone delivery in *Plethodon spp*., the number and diversity of PMF genes rapidly expanded with the mental gland expressing >30 rapidly evolving isoforms (Wilburn et al., 2012). Given the extraordinary molecular sensitivity and specificity of ligand binding to olfactory receptors (Leinders-Zufall et al., 2009), we hypothesized that transition from transdermal to olfactory delivery reduced the minimum concentration of PMF necessary to stimulate female receptors and expressing more PMF isoforms maximizes the chances of male reproductive success (Wilburn et al., 2014b). By contrast, the expression of a single highly abundant spPMF isoform by *P. shermani* sperm may be indicative of stoichiometric binding to an abundant egg receptor.

**Figure 6.**
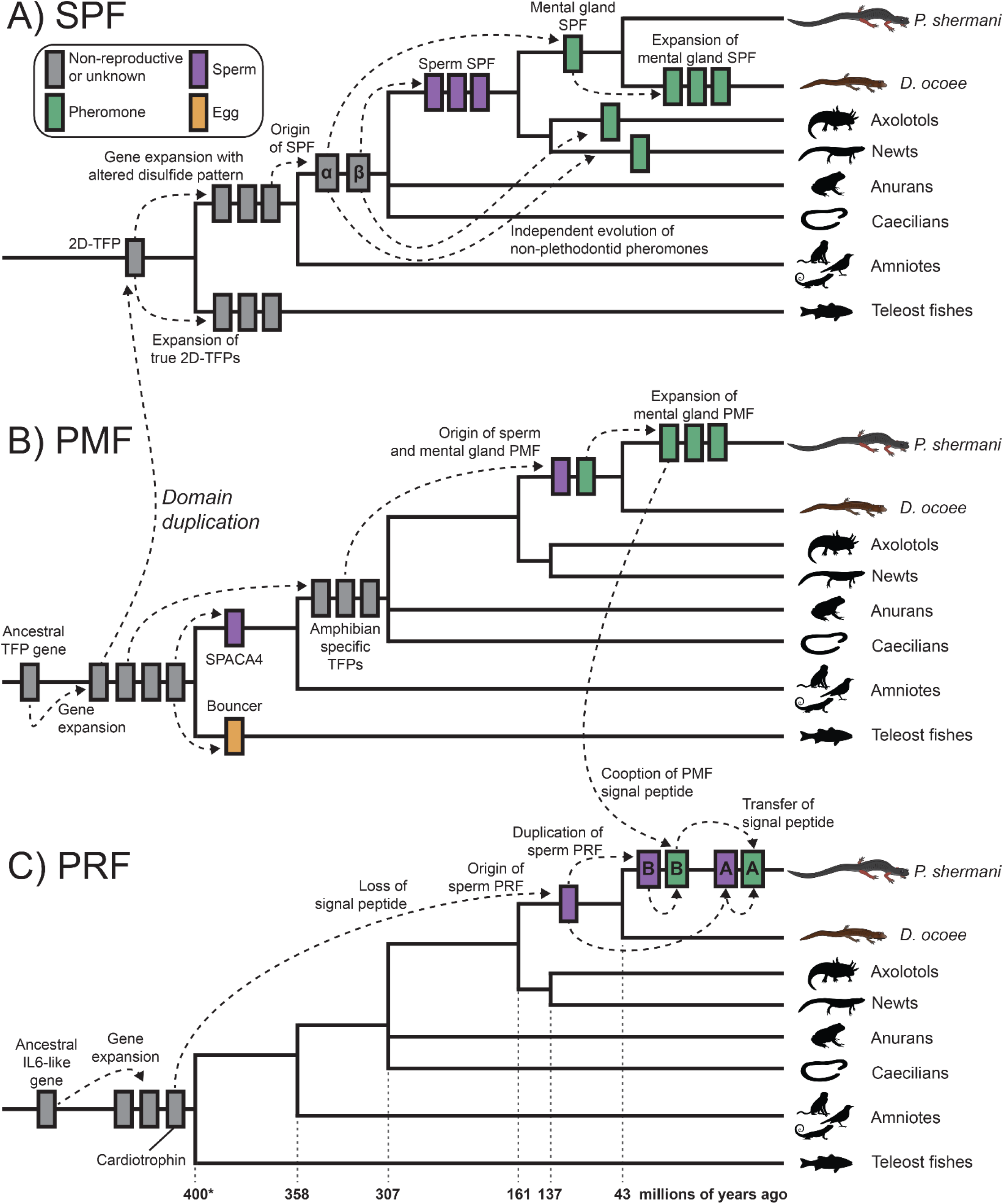
Proposed model of cytokine and TFP evolution generating both plethodontid pheromones and sperm proteins. (A) An ancestral tandem duplication of a 10C-TFP protein produced the first 2D-TFP that subsequently expanded into independent families, with teleost fish having 2D-TFPs that retained the canonical disulfide patterns, and tetrapods having a separate family with a different disulfide pattern (purple in Figure 4). Further duplication of these tetrapod proteins resulted in the SPFα and SPFβ families within amphibians that independently gave rise to multiple descendants, including plethodontid pheromones from SPFα and spSPF from SPFβ. Within plethodontids, further expansion of pheromone SPFs occurred in *D. ocoee* relative to *P. shermani*. (B) A progenitor 8C-TFP gene expanded at the base of vertebrates giving rise to several families, including the 10C-TFP from which SPF is originally derived. Another 10C-TFP led to the evolution of SPACA4 in tetrapod sperm and Bouncer in fish oocytes. Continued evolution of an 8C-TFP family led to the evolution of PMF in plethodontid salamanders, likely through an amphibian-specific intermediate, with sperm and mental gland paralogs evolving nearly simultaneously. Following transition from transdermal to olfactory delivery, there was a large expansion of PMF genes in *P. shermani* relative to *D. ocoee*. (C) PRF is distantly related to IL-6 cytokines and expanded into several related genes early in vertebrate evolution, including a cardiotrophin-like gene. Within plethodontid salamanders, a descendent of the cardiotrophins duplicated and lost its signal peptide to yield spPRF. Continued expansion of spPRF genes in *Plethodon spp*. resulted in spPRF-A and B that independently duplicated into PRF-A and B, respectively. PRF-A and B seemingly reacquired extracellular expression through recombination with a PMF gene. Based on the sequence alignment, we propose that PRF-B first co-opted the PMF signal peptide (and partial 5’ UTR sequence), with PRF-A acquiring its signal peptide by recombination with PRF-B.

Of the three major pheromone families, PRF has perhaps the most convolved evolutionary history: a secreted cardiotrophin-like ancestor lost its signal peptide to become a likely cytoplasmic sperm protein, followed by a gene duplication of this spPRF recombining with PMF to reacquire secretion as a pheromone. It is especially remarkable that this signal peptide reacquisition for pheromone PRF occurred twice, with PRF-A and PRF-B arising independently from spPRF-A and spPRF-B, respectively. Based on 5’ UTR and signal peptide sequences, we hypothesize this occurred through a two-step duplication process where PRF-B acquired its signal peptide from PMF class III, and then PRF-A acquired its signal peptide from PRF-B. While PMF and other TFPs are highly prone to gene duplication, it is conspicuous that the PRF signal peptide arose from a highly expressed gene in the same tissue and may suggest an important role of transcriptionally active genes being more prone to gene duplication because of chromatin structure. This would be especially relevant for highly expressed gametic genes where duplications could fix in the germline and be more exposed to natural or sexual selection. Abundant expression of all three pheromones in the plethodontid sperm proteome (although at lower levels than the sperm paralogs) may indicate leaky expression of androgen responsive genes during sperm maturation, which could expose these pheromone loci to similar modes of gene evolution as their sperm-specific paralogs. This complex interplay between genes expressed by male secondary sex characteristics can likely help explain both the extraordinary number and the rapid evolution of plethodontid pheromone proteins. In summary, our transcriptomic and proteomic analyses of plethodontid gametic proteins highlight the novelty of reproductive proteins present in poorly characterized species, such as salamanders, and how new fertilization proteins may evolve to facilitate species-specific fertilization in vertebrates.

## Author contributions

DBW, RCF, and PWF contributed to the conception and design of the study. DBW, CLK, RCF, and PWF performed field collections, animal care, staged courtship trials, and dissection for tissue collection. DBW prepared transcriptomic libraries and proteomic samples. BCS performed all mass spectrometry experiments. DBW and BCS analyzed the data. DBW drafted the manuscript with revisions from BCS and RCF. All authors read and approved the submitted version.

## Acknowledgements

We would like to thank Dr. Willie Swanson for feedback on molecular evolutionary analyses performed in this study, as well as Dr. Jim Costa and the staff of Biological Station for their valuable assistance and support that enable the salamander experiments in this study.

## Data Availaility Statement

Pacbio Iso-Seq data has been deposited in the NCBI Sequence Read Archive (BioProject ID PRJNA785352). Mass spectrometry data have been deposited on the ProteomeXchange Consortium via the PRIDE partner repository with the dataset identifier PXD030143. The presented pylogenetic trees and their underlying sequence alignments are provided in the supplementary material.

