## Supplementary Figures for "Recurrent co-option and recombination of cytokine and three finger proteins in multiple reproductive tissues throughout salamander evolution"

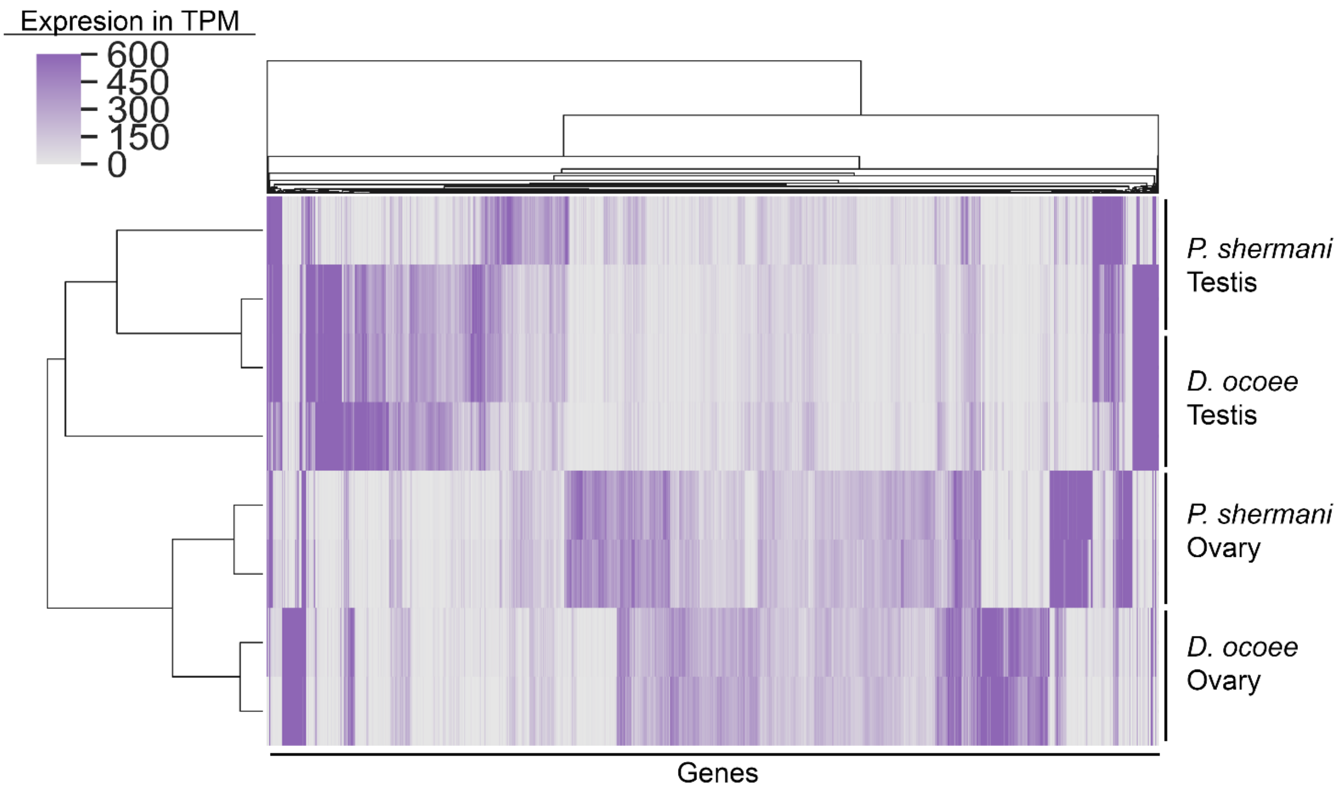


**Supplementary Figure 1.** Heat map of gene expression for the gonad transcriptome that includes testis and ovary of both *P. shermani* and *D. ocoee* in biological duplicate, with genes and samples arranged based on hierarchical clustering. Testis gene expression is highly variable both within and between species compared to ovary expression.

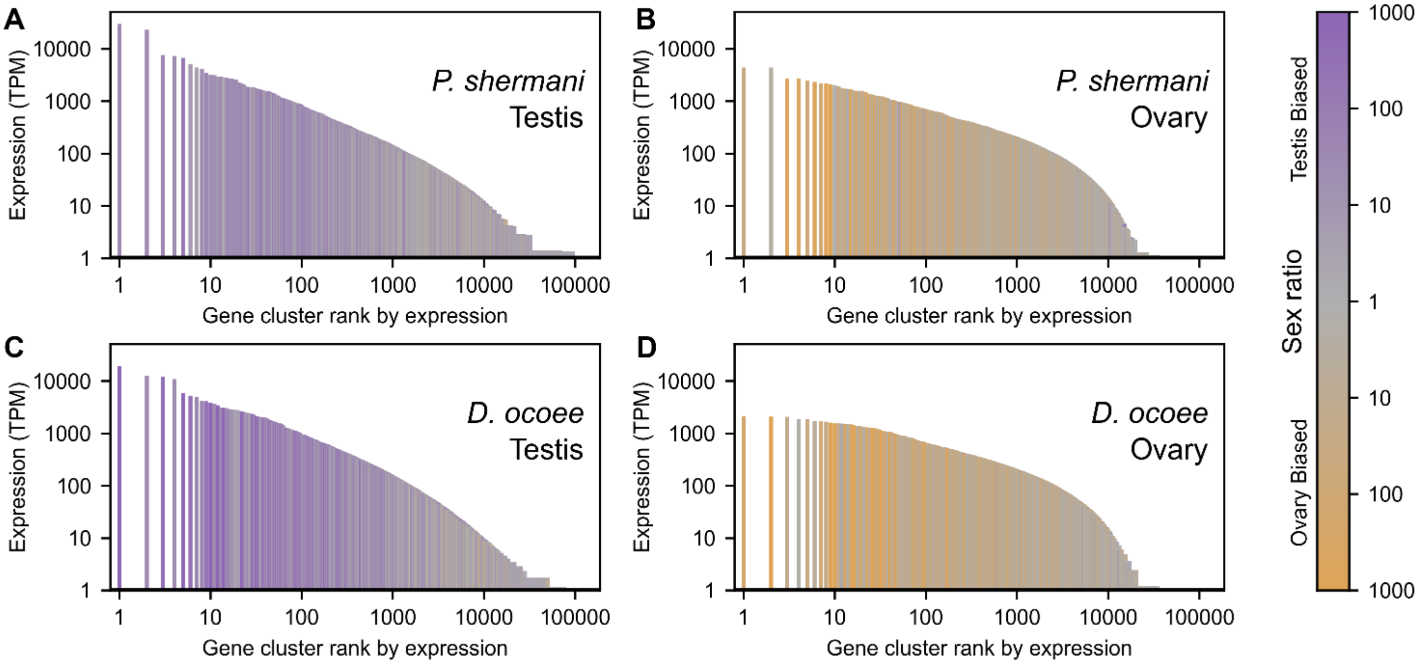


**Supplementary Figure 2.** Mean transcript abundance in TPM (plotted in log scale) for testis and ovary from both *P. shermani* and *D. ocoee* organized by expression rank (in log scale) for each species-tissue combination, with bars shaded according sex ratio bias. The most highly expressed testis genes are ~10x more abundant than the most highly expressed ovary genes. Testis also expresses many more genes in total compared to ovary such that ovary expression is generally more balanced compared extreme properties in testis expression.
